# High-risk Autism Spectrum Disorder Utah pedigrees: a novel Shared Genomic Segments analysis

**DOI:** 10.1101/134957

**Authors:** Todd M Darlington, Deborah Bilder, Jubel Morgan, Leslie Jerominski, Venkatesh Rajamanickam, Rob Sargent, Nicola J Camp, Hilary H Coon

**Affiliations:** Department of Psychiatry, University of Utah, Salt Lake City, Utah 84108; Department of Internal Medicine, Division of Cancer Control and Population Sciences, University of Utah, Salt Lake City, Utah 84108

## Abstract

Progress in gene discovery for Autism Spectrum Disorder (ASD) has been rapid over the past decade, with major successes in validation of risk of predominantly rare, penetrant, *de novo* and inherited mutations in over 100 genes (de Rubies et al., 2015; Sanders et al., 2015). However, the majority of individuals with ASD diagnoses do not carry a rare, penetrant genetic risk factor. In fact, recent estimates suggest that most of the genetic liability of ASD is due to as yet undiscovered common, less penetrant inherited variation (Gaugler et al., 2014) which is much more difficult to detect. The study of extended, high-risk families adds significant information in our search for these common inherited risk factors. Here, we present results of a new, powerful pedigree analysis method (Shared Genomic Segments—SGS) on three large families from the Utah Autism Research Program. The method improves upon previous methods by allowing for within-family heterogeneity, and identifying exact region boundaries and subsets of cases who share for targeted follow-up analyses. Our SGS analyses identified one genome-wide significant shared segment on chromosome 17 (q21.32, p=1.47x10^-8^). Additional regions with suggestive evidence were identified on chromosomes 3, 4, 6, 8, 11, 13, 14, 15, and 18. Several of these segments showed evidence of sharing across families. Genes of interest in these regions include *ATP8A1*, *DOCK3*, *CACNA2D2*, *ITGB3*, *AMBRA1*, *FOLH1*, *DGKZ*, *MTHFS*, *ARNT2*, *BTN2A2*, *BTN3A1*, *BTN3A3*, *BTN2A1*, and *BTN1A1*. We are exploring multiple other lines of evidence to follow up these implicated regions and genes.

## Introduction

Autism Spectrum Disorder (ASD) affects an estimated 1 in 68 children (Centers for Disease Control and Prevention, 2016), and imposes an enormous psychological and economic burden on affected individuals, their families, and society [1]. Genetic risk factors are important in the etiology of ASD, a conclusion strongly supported by recent population-based twin studies [2,3], and an additional recent twin meta-analysis [4]. These results suggest not only strong genetic effects, but also complex inheritance, with heterogeneity between individuals, in addition to the likely involvement of multiple interacting genes within individuals. Indeed, recent genetic discovery studies have identified numerous inherited and *de novo* variants, and place the estimates for numbers of causal variants anywhere from several hundred to over a thousand [5–7].

Given this complex landscape, studies of large extended pedigrees provide an opportunity to add value. Allelic and locus heterogeneity are reduced and the ability to detect familial variants is enhanced [8,9]. Familial sharing of segments of the genome have the potential to provide important prioritization for the analysis of sequence variation, and may also add essential information about transmission, penetrance, and specificity provided by family relatives. Importantly, extended pedigrees offer sufficient statistical power to detect familial variants, often in a single pedigree, through the efficient study of relatively few individuals, although care must be taken to determine variants present in the pedigree relatives contributing to familial sharing to focus subsequent follow-up work. In addition, evidence is emerging that some variants contribute to phenotypes related, but not always specific to autism, such as intellectual disability [10–12]. The analysis of family members will also allow investigation of potential associations with phenotypes related to ASD in clinically affected and unaffected relatives.

The Utah Autism Research Program has ascertained a resource of high-risk extended families using the Utah Population Database (UPDB), a rich resource of health data and genealogical information for over eight million individuals who include descendants of the nineteenth century Utah pioneers [13]. Previous studies have shown low rates of inbreeding within the UPDB [14,15], and the population is primarily of Northern European ancestry [16]. We used the genealogical data within the UPDB resources to identify extended pedigrees with significantly increased incidence of ASD. This analysis focuses on the identification of segments of the genome shared among affected individuals in three of these extended families, and shared by few of their unaffected relatives, followed by a presentation of possible causal exome sequence variants within these segments. While we have previously explored these extended families using linkage methods [17], here we have employed the novel method Shared Genomic Segments (SGS) [18]. This method has the advantage of identifying which pedigree members share which genomic segments, allowing for more accurate subsequent follow-up identification of possible variation explaining the pedigree sharing in sequence data. An ability to focus on shared regions in specific individuals will be of particular importance as we map more common variants that may exist outside the exome.

## Materials and Methods

### Subjects

Subjects were selected from three of the largest high-risk extended families in a previously described research sample [17,19]. Extended families in our study were identified or confirmed using the UPDB. Many additional distant family relationships between individuals with ASD that were not known to the subjects or their immediate families were identified. Each of the three large families in this study has multiple generations connecting many related nuclear families (see Figure 1). We analyzed data from a total of 24 individuals with ASD (8 each from families 10001, 25002, and 8002). In family 10001, there is one monozygotic affected twin pair; one member of the pair (64049) was randomly selected for this analysis. Additionally, at the data QC phase of the project, one individual with ASD in family 25002 was found to have a chromosome anomaly, and was therefore omitted from further analyses (83681), bringing the number of affected cases in 25002 in the final analyses to 7. We also studied 23 unaffected siblings in these families (8 in 10001, 8 in 25002, and 7 in 8002). Table 1 shows the characteristics of these individuals, and they are pictured in Figure 1. This study has ongoing approval from the University of Utah Institutional Review Board (approval IRB_00006042) and the Resource for Genetic and Epidemiologic (RGE) committee which governs the use of UPDB data. Written consent was obtained for all study subjects.

**Figure 1.**
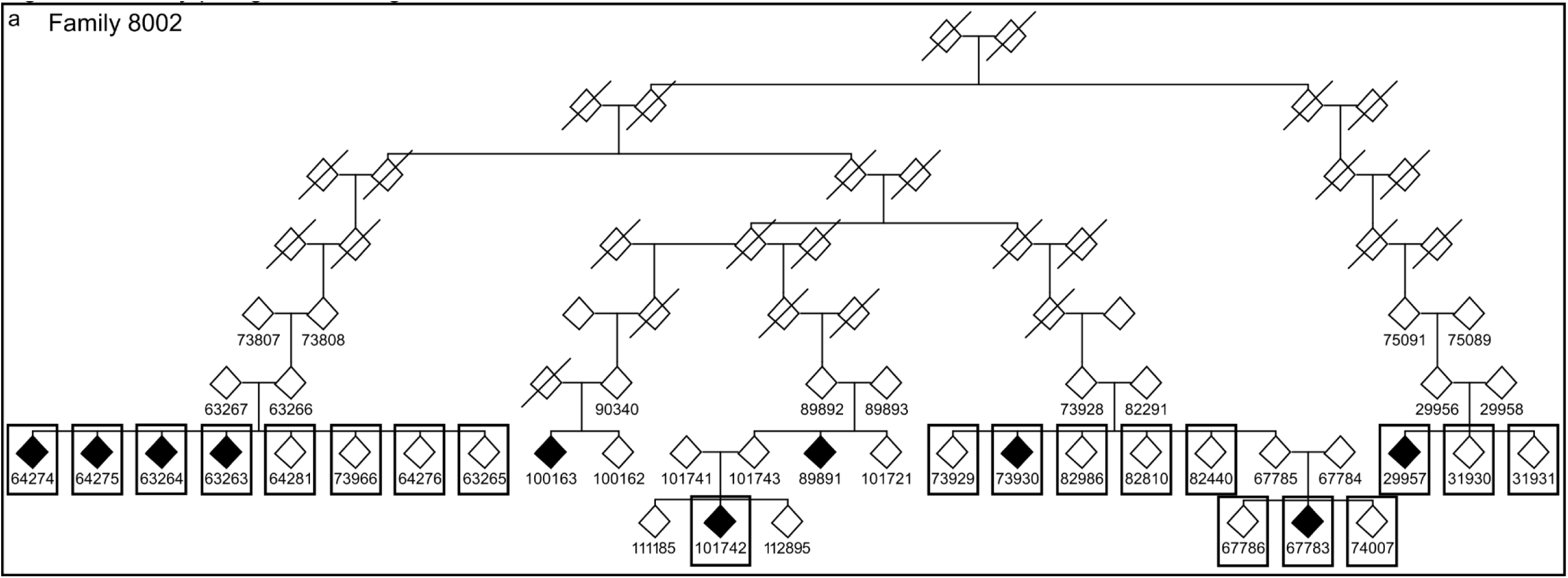
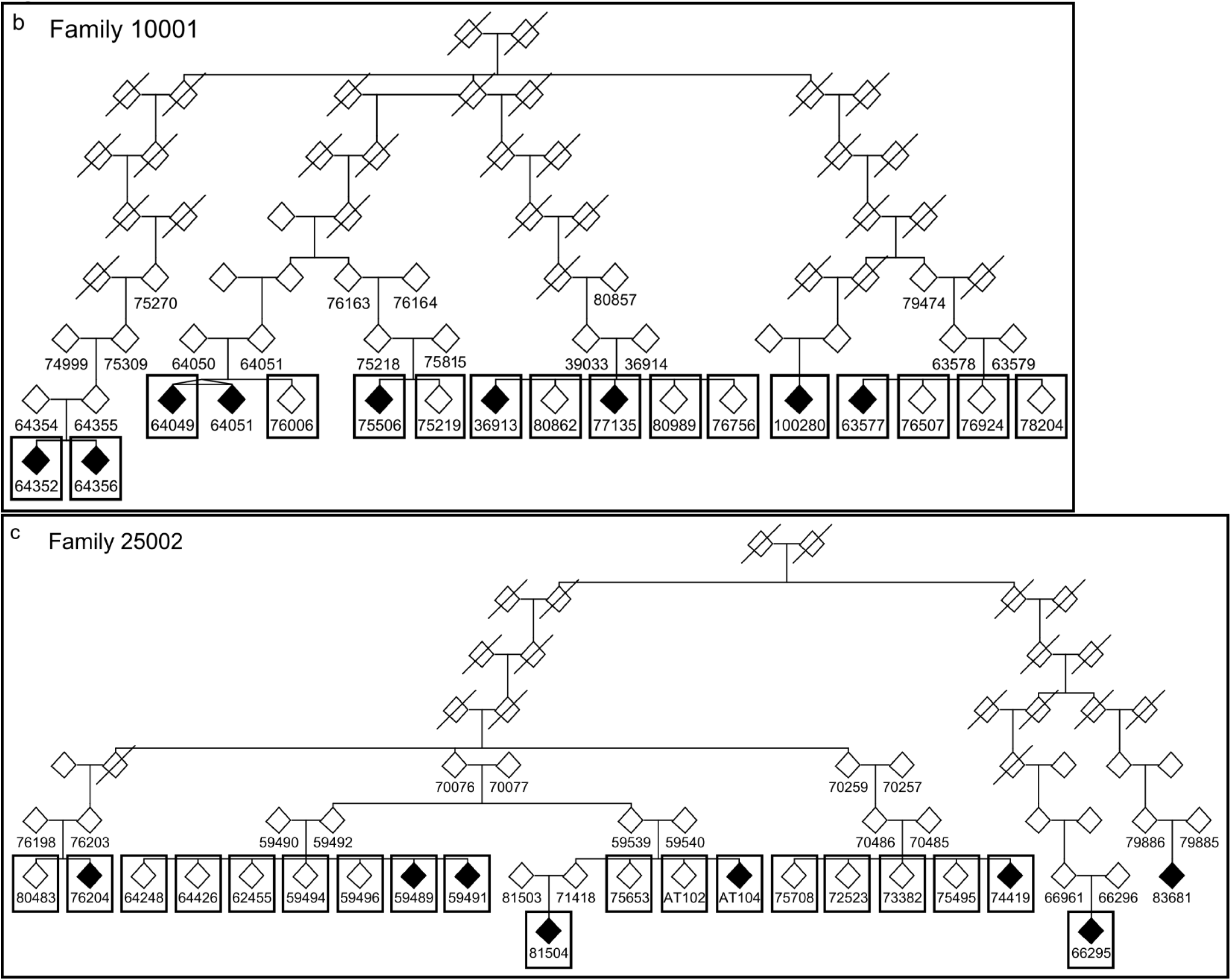
Family pedigree drawings. Gender has been disguised, and sibling order has not been maintained in order to protect family privacy. Shaded symbols represent individuals with ASD. All family members with a numeric ID have available DNA. Boxed cases and unaffected siblings were used for SGS analyses. OmniExpress genotype data from family 8002 members 89891 and 100163 were not available at time of analysis (a). Family 10001 members 64049 and 64052 are monozygotic twins; 64049 was selected randomly for analysis (b). Family 25002 member 83681 was found during quality control to have a chromosomal anomaly and therefore the containing branch was not used for analysis (c).

**Table 1.**
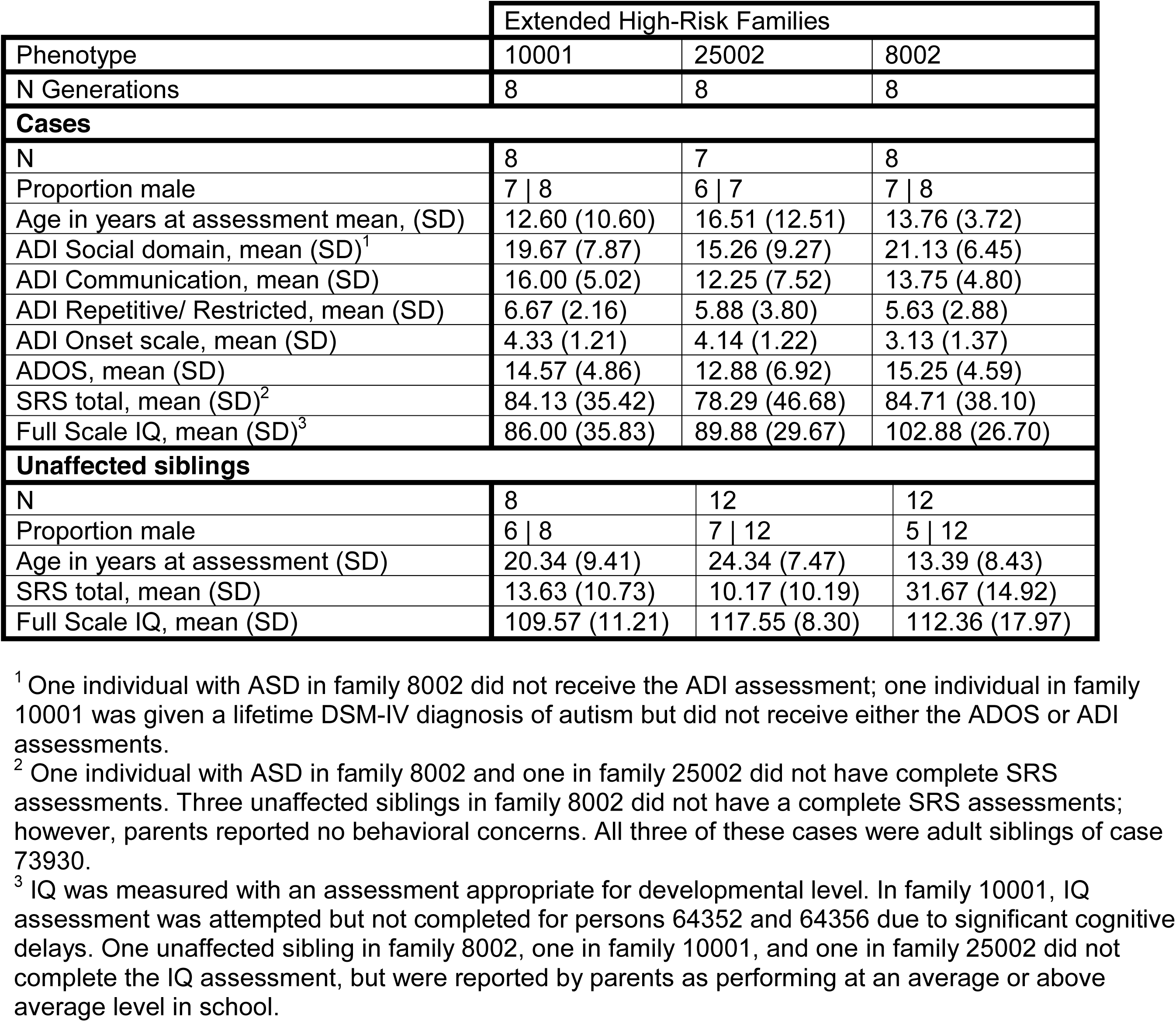
Description of individuals with ASD and unaffected siblings used for Shared Genomic Segments (SGS) analyses in three extended families

### Phenotyping

Family members were requested to complete questionnaires and in-person testing. ASD was diagnosed with the Autism Diagnostic Interview-Revised (ADI-R) [20] and the Autism Diagnostic Observation Schedule (ADOS) [21], for all but two affected participants. One of these was given the ADOS only; the other received a lifetime DSM-IV diagnosis of autism by a clinical expert. Participants with ASD had no other reported major medical conditions or evidence of brain injury. One affected individual in family 8002 and one in family 10001 were non-verbal; all other affected individuals were verbal.

All but two affected participants had data available for the Social Responsiveness Scale, a quantitative measure of ASD characteristics ranging from significant impairment to above average social abilities [22,23]. SRS data were also obtained for all unaffected siblings with the exception of three adult unaffected siblings of case 73930 in family 8002. The parents reported no behavioral concerns for these siblings.

IQ was assessed using measurements appropriate for age and verbal ability, as described in Coon et al (2010) [17]. All affected cases had IQ estimates except two cases in family 10001; these two subjects were not testable due to intellectual disability. For the two unaffected siblings missing IQ in this study, parents reported average or above average school performance. Additionally, other self-reported medical history data were collected on most participants.

### Genotyping

Subjects were genotyped on the Illumina OmniExpress v1.0 platform using protocols and standards as set by Illumina. Raw data were output as Illumina forward orientation. To accurately compare our data with the publicly available control set from the 1000 Genomes Project (1KGP) [24], our data was converted to 1KGP forward orientation. A reference panel of 152 subjects with data from both the Illumina OmniExpress v1.0 platform and sequencing in 1KGP were used to determine a translation between the two platforms. A total of 25,166 variants were discarded due to inconsistency between the platforms. Only bi-allelic, single nucleotide variants (SNVs) were used.

### Quality control - Plink

Quality control was performed using PLINK [25] on the whole set of pedigrees combined with 168 control subjects of European descent from the 1KGP. First, 49 variants were removed with less than 95% call rate. No subjects were removed for genotype rates less than 90%. Next, sex checks were performed, resulting in one subject removed for a chromosomal anomaly. Relatedness estimates confirmed relationship status between pedigree members. A total of 12,688 variants were discarded due to deviation from Hardy-Weinberg equilibrium. Finally, 8,402 variants were removed for likely genotype errors, identified by Mendelian misinheritance, and 8,100 uninformative, monomorphic markers were removed prior to analysis.

### Shared Genomic Segments

Plink format files were converted to SGS format using software available through http://genepi.med.utah.edu/∼alun/software/index.html. Genetic distance was interpolated using the Rutgers map (v3) [26]. Variants from 168 northern European 1KGP controls were used to construct a graphical model of linkage disequilibrium, used for simulating founder genotypes.

A pedigree SGS analysis considers sampled, distantly related cases (without genotyping from connecting relatives) and poses the question as to whether the length of consecutively shared loci (identified as identical-by-state, or IBS) among affected cases is longer than expected by chance [18]. IBS is established by determining if alleles at sequential loci are the same (phase is ignored). IBS is not the same as identity-by-descent (IBD; the same inherited segment from a common ancestor) which is our true interest; however, if the length of SGS shared IBS is significantly longer than expected by chance (given the known pedigree relationships) then IBD is implied. Observed sharing statistics (length of sharing) for a given pedigree are compared to simulated (“null”) shared statistics for an identical pedigree structure to determine empirical probabilities. Simulations involve assigning founder haplotypes (based on the graphical model of linkage disequilibrium) followed by simulating Mendelian inheritance (with recombination likelihoods derived from Rutgers map) to all pedigree members (“gene-dropping”). For the length of a simulated segment to surpass an observed segment, the simulated shared segment must span the entire observed segment. For each pedigree, all possible combinations of affected cases were assessed for sharing. The total number of simulations to determine empirical significance for each subset of cases and for each chromosome ranged from 300,000 to 200,000,000. After a minimum of 300,000 simulations, no further simulations were performed on any particular subset where the most significant segment was observed at least three times in simulated data.

Thresholds for significance were determined using a modification of the technique previously described by Lander and Kruglyak (1995) for assessing significance in linkage analysis [27]. The modification incorporates extreme value theory and the law of large deviations in order to account for multiple testing (See supplemental methods). Briefly, empirical p-values for each marker were optimized by selecting the lowest p-value at each marker across all subsets, and reduced such that any shared segment in the optimized set was represented only once. The resulting distribution of p-values was used for threshold determination as described in supplemental methods. Additionally, to identify common segments shared across multiple families, the optimized p-values from each family were combined across families using Fisher’s method in R (metap, v0.7). There were two comparisons. First, we compared all families by combining optimized p-values across all three families. Second, we compared pairs of families by combining optimized p-values across each pair, then re-optimized at each marker using the combined p-values from each of the three pair combinations (8002-10001, 8002-25002, 10001-25002). Since these two comparisons are not independent, the thresholds were adjusted accordingly.

Although SGS infers IBD in shared segments, it only directly measures IBS. Therefore, for any significant SGS regions, the actual IBD segments are likely flanked by segments of random IBS, i.e. the actual interesting segment may be smaller than the SGS region identified. To address this, we utilized the presence of multiple affected siblings in each of our families to identify the most likely IBD segment, as follows. Consider a hypothetical family consisting of three affected cases. Case1 and Case2 are siblings, and Case3 is distantly related. Case2 and Case3 share a segment spanning markers A, B, C, D, and E. However, when considering Case1, Case2, and Case3, the shared segment only spans A, B, and C, so therefore we infer that markers D and E were likely random IBS between the two distant cousins, and the actual IBD region is A, B, and C.

To assess preferential segregation of identified shared segments to ASD cases, significant segments were re-analyzed including all possible combinations also of unaffected siblings of cases, who serve as within-family controls. We assume variants associated with ASD risk may have reduced penetrance and/or may affect sub-clinical aspects of ASD that can occur in unaffected siblings; therefore, some degree of segregation to these siblings is not unexpected. However, excessive segregation to unaffected siblings may indicate a false positive result. Although we prioritized findings where less than one half of within-family controls shared the same segment, all segments with evidence of IBD are reported.

### Online bioinformatics for segment follow-up

Base pair boundaries for each shared segment were defined as: previous unshared marker base position + 1bp before the shared segment to the first unshared marker base position - 1bp after shared segment. Genes within shared segments were identified using the Table Browser tool [28] from the UCSC Genome Browser [29,30]. Known ASD risk genes were identified using ASD@Princeton [31], a new, validated, web-based tool developed using machine-learning techniques based on known ASD genes and a human brain-specific gene network.

### Plotting

Plots and visualizations were generated using the UCSC Genome Browser, R v3.2.2 [32] and ggplot2 v1.0.1 [33], Progeny v9.5.0.2 pedigree drawing software, and the Genome Decoration Page tool (http://www.ncbi.nlm.nih.gov/genome/tools/gdp/).

## Results

### Shared Genomic Segments

Significance thresholds for families and shared segments meeting those thresholds in each family are shown in Table 2 and Figure 2. One segment on chromosome 17 from family 10001 reached genome-wide significant (chr17q21.32, p=1.47x10^-8^), with only a single unaffected sibling (76756) also sharing. Several segments in each family exceeded genome-wide suggestive thresholds. In family 8002, three shared segments reached the suggestive threshold, two on chromosome 4 and one on chromosome 13. Segregation of segments ranged from 3-9 internal controls out of 12 total. In family 10001, three segments reached the suggestive threshold, one each on chromosomes 3, 6, 14. Segregation ranged from 3-7 out of 8 total internal controls. In family 25002, two segments, one each on chromosome 11 and 15, reached the suggestive threshold with 3-7 out of 12 internal controls also sharing. Shared segments for each family are shown in Figure 3, along with genes in each segment. Genes were only included figure if they resided in the shared segment of interest (Figure 3). Only segments with genes are shown. Specific subset details are given in Supplemental Material.

**Table 2.** Shared genomic segments with at least genome-wide suggestive evidence in each family.

| **Family** | **Locus** | **Cytoband** | **p-value<sup>a</sup>** | **Cases<sup>b</sup>** | **Controls<sup>c</sup>** |
| --- | --- | --- | --- | --- | --- |
| **8002** | chr4:42,646,678-43,197,961 | 4p13 | 6.64x10<sup>-7</sup> | 7 | 3/12 |
| **8002** | chr4:157,340,211-157,601,192 | 4q32.1 | 1.85x10<sup>-7</sup> | 8 | 6/12 |
| **8002** | chr13:23,586,367-23,641,247 | 13q12.12 | 6.53x10<sup>-7</sup> | 7 | 9/12 |
| **10001** | chr3:50,421,082-51,927,597 | 3p21.31-p21.2 | 5.51x10<sup>-8</sup> | 8 | 3/8 |
| **10001** | chr3:52,348,365-53,160,359 | 3p21.1 | 5.51x10<sup>-8</sup> | 8 | 4/8 |
| **10001** | chr6:130,537,006-131,438,062 | 6q23.1-q23.2 | 4.91x10<sup>-8</sup> | 7 | 3/8 |
| **10001** | chr14:59,505,554-59,585,931 | 14q23.1 | 1.94x10<sup>-7</sup> | 8 | 4/8 |
| **10001** | chr17:45,300,219-46,574,490 | 17q21.32 | 1.47x10<sup>-8*</sup> | 7 | 1/8 |
| **25002** | chr11:46,309,969-47,394,304 | 11p11.2 | 1.20x10<sup>-6</sup> | 7 | 6/12 |
| **25002** | chr11:47,449,073-50,519,186 | 11p11.2-p11.12 | 1.20x10<sup>-6</sup> | 7 | 7/12 |
| **25002** | chr15:79,604,783-80,724,488 | 15q25.1 | 1.47x10<sup>-6</sup> | 6 | 3/12 |

<sup>a</sup>Significant and suggestive thresholds for family 8002 (9.90x10<sup>-8</sup> and 1.05x10<sup>-6</sup>); 10001 (1.92x10<sup>-8</sup> and 2.94x10<sup>-7</sup>); 25002 (2.20x10<sup>-7</sup> and 2.44x10<sup>-6</sup>)
<sup>a</sup>Number of affected cases sharing this segment
<sup>b</sup>Number of internal controls that share this segment with affected cases, shown with total number of possible internal controls
\*Genome-wide significant

**Figure 2.**
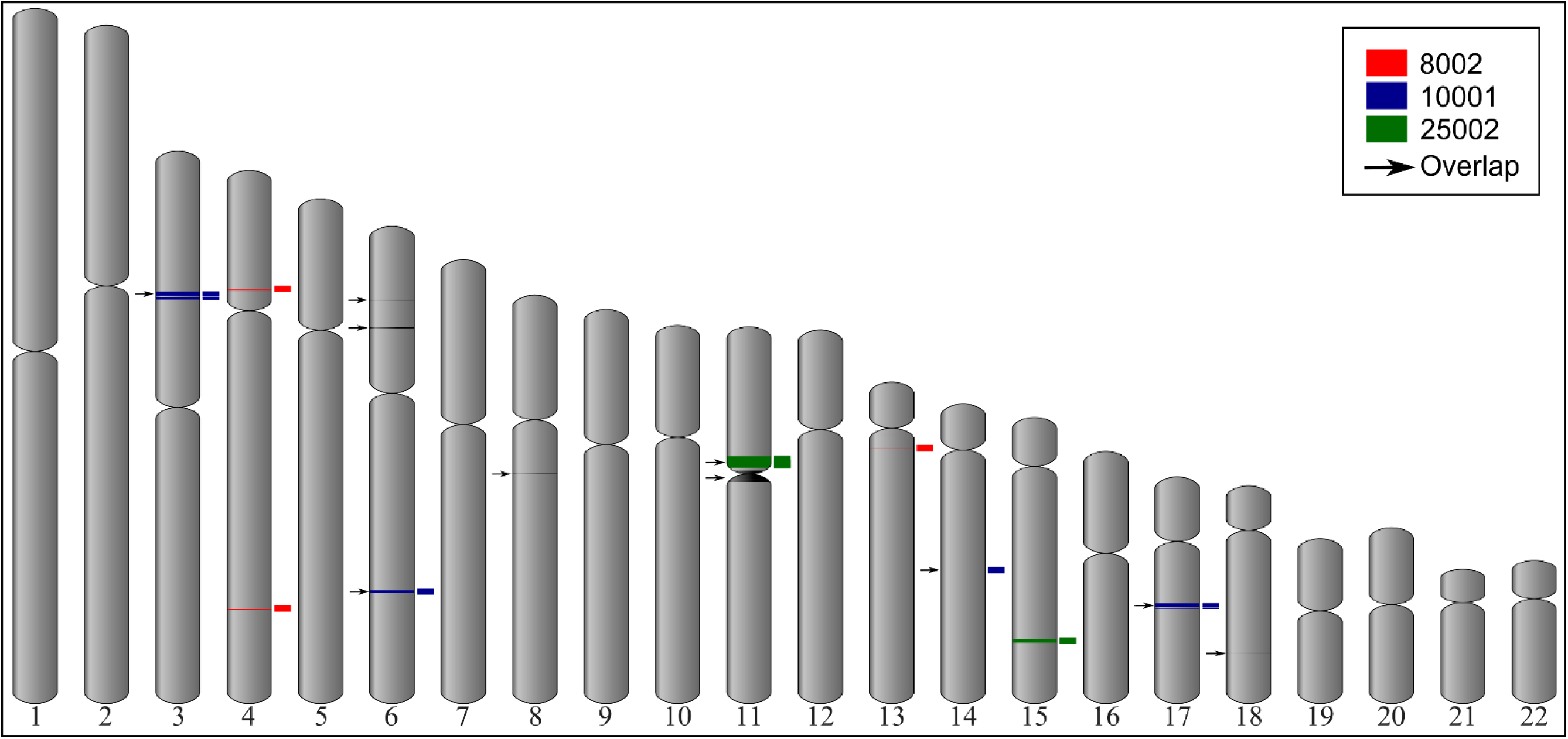
Shared genomic segments meeting significance thresholds. Shared genomic segments for each family and overlapping segments across multiple families are shown for each autosomal chromosome. Chromosome number is shown below each chromosome.

**Figure 3.**
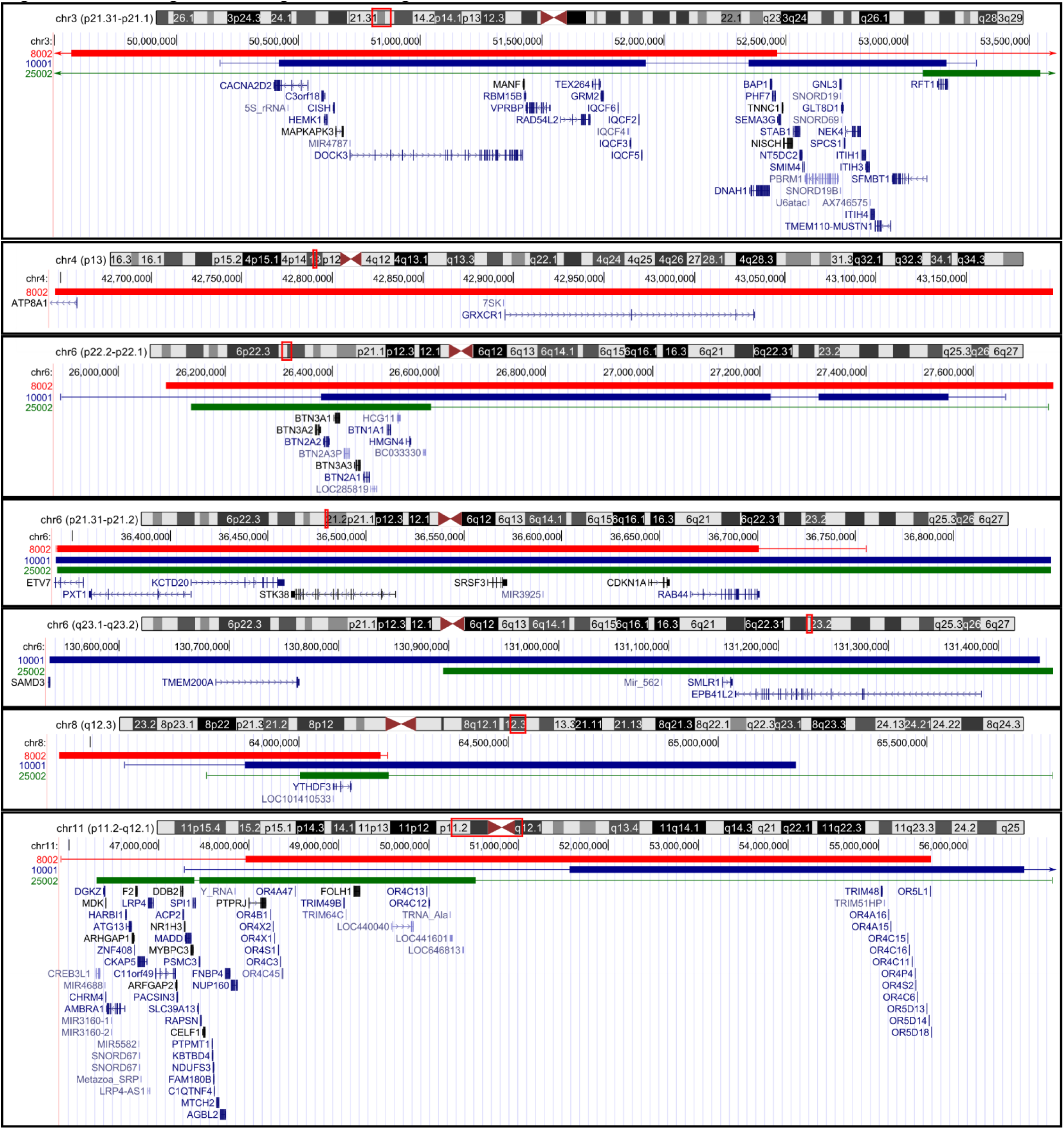
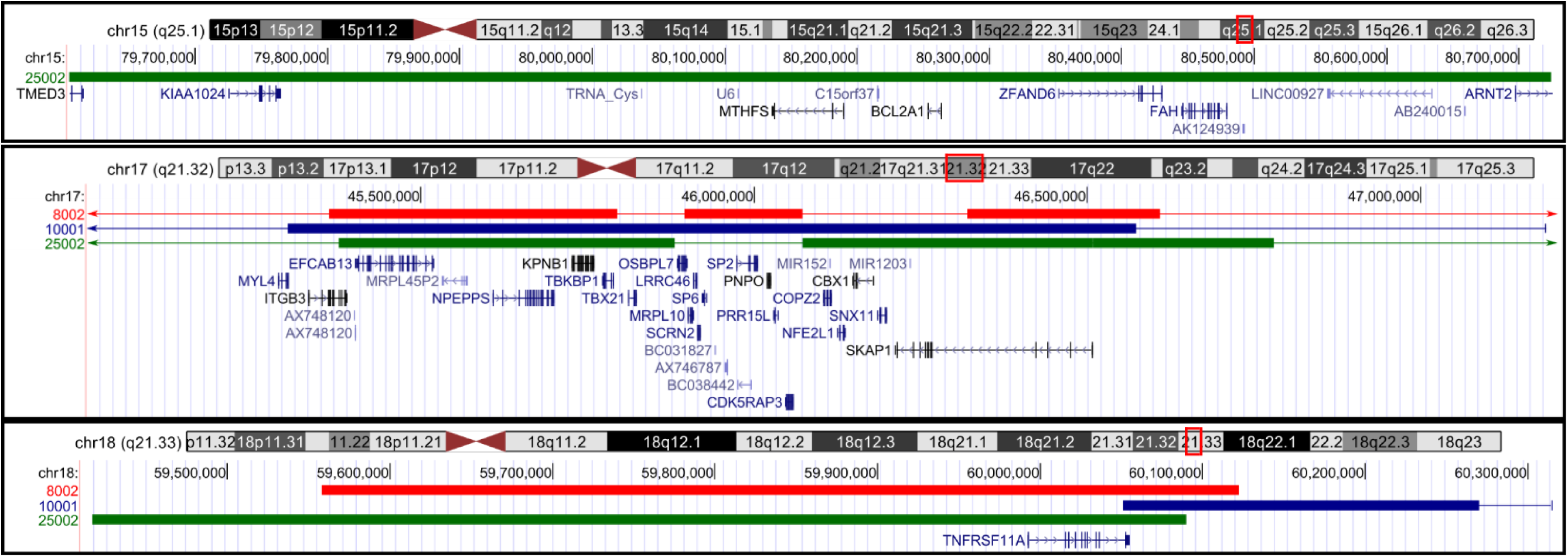
Shared genomic segments with genes. Figure 3 shows segments of the genome shared between affected cases in each family that meet significance thresholds. Chromosome segments are denoted by red boxes on each chromosome ideogram. Shared segments from family 8002 (red), 10001 (blue), 25002 (green) are shown below each ideogram. Thick portions of each family bar represent estimated actual shared segment. Thin bars for each family, if present, represent entire segment originally identified by SGS. Actual shared segments were determined by including any missing siblings from the original subset. Genes are displayed below shared segments.

### Overlapping segments from multiple families

Significance thresholds for segments identified in multiple families and shared segments meeting those thresholds in each family are shown in Table 3 and Figure 2. Several segments met the suggestive threshold. When considering multiple families, an additional 13 regions are identified. These segments are shown in Figure 3 along with the genes in each segment. Again, genes were included in the figure if they resided in the shared segment of interest. Only segments with genes are shown.

**Table 3.** Overlapping shared segments with at least genome-wide suggestive evidence across families.

| Families | Locus | Cytoband | p-value<sup>a</sup> | #Cases;#Controls<sup>b</sup> |  |  |
| --- | --- | --- | --- | --- | --- | --- |
|  |  |  |  | 8002<br>8;12 | 10001<br>8;8 | 25002<br>7;12 |
| 8002;10001 | chr3:50,421,082-51,927,597 | 3p21.31-p21.2 | 1.20x10<sup>-10</sup> | 5;6 | 8;3 |  |
| 8002;10001 | chr3:52,348,365-52,467,323 | 3p21.1 | 1.20x10<sup>-10</sup> | 5;6 | 8;4 |  |
| 8002;10001 | chr11:51,566,910-55,584,108 | 11p11.12-q11 | 2.19x10<sup>-10</sup> | 6;7 | 6;4 |  |
| 8002;10001 | chr14:59,505,554-59,521,231 | 14q23.1 | 3.70x10<sup>-10</sup> | 7;9 | 8;4 |  |
| 8002;25002 | chr11:47,969,153-50,519,186 | 11p11.2-p11.12 | 4.53x10<sup>-10</sup> | 6;8 |  | 7;7 |
| 10001;25002 | chr6:130,894,603-131,438,062 | 6q23.1-q23.2 | 2.54x10<sup>-10</sup> |  | 7;3 | 3;3 |
| All | chr6:26,378,682-26,584,667 | 6p22.2 | 2.59x10<sup>-12</sup> | 3;3 | 7;6 | 6;10 |
| All | chr6:36,342,182-36,700,832 | 6p21.31-p21.2 | 4.57x10<sup>-12</sup> | 8;8 | 6;4 | 2;0 |
| All | chr8:64,001,937-64,194,109 | 8q12.3 | 4.82x10<sup>-12</sup> | 7;0 | 5;1 | 5;5 |
| All | chr17:45,377,578-45,794,705 | 17q21.32 | 7.05x10<sup>-13</sup> | 8;6 | 7;1 | 6;0 |
| All | chr17:46,319,649-46,509,446 | 17q21.32 | 7.05x10<sup>-13</sup> | 8;7 | 7;1 | 6;0 |
| All | chr17:46,509,448-46,574,490 | 17q21.32 | 7.05x10<sup>-13</sup> | 8;7 | 7;1 | 6;0 |
| All | chr18:60,050,642-60,090,078 | 18q21.33 | 6.48x10<sup>-12</sup> | 6;4 | 6;2 | 4;2 |

<sup>a</sup>Significant and suggestive thresholds for pairs of families (2.30x10<sup>-11</sup> and 4.53x10<sup>-10</sup>); all three families (2.00x10<sup>-13</sup> and 7.34x10<sup>-12</sup>)
<sup>b</sup>Number of affected cases and unaffected siblings sharing the respective segment in each family. The total number of affected cases and unaffected siblings per family is in the header line.

## Discussion

This analysis has revealed several familial shared genomic segments that may harbor variants contributing to Autism Spectrum Disorder risk in specific cases in our study. From within-family analyses, these segments are on chromosomes 3, 4, 6, 11, 13, 14, 15, and 17. Several of these segments also showed evidence of sharing between families. Additional between-family shared segments were identified on chromosomes 6, 8, 11, and 18.

There were several genes in these segments that have been previously implicated in autism spectrum disorders and multiple others that have been associated with autism-like phenotypes. Specific genes of interest include *ATP8A1* in family 8002; *DOCK3*, *CACNA2D2*, and *ITGB3* in family 10001; *AMBRA1*, *FOLH1*, *DGKZ*, *MTHFS*, and *ARNT2* in family 25002; and several butyrophilin subunits (*BTN2A2*, *BTN3A1*, *BTN3A3*, *BTN2A1*, and *BTN1A1*) in segments shared across multiple families.

In family 8002, seven affected members share the segment on chromosome 4p13 containing ATPase phospholipid transporting 8 A1 (*ATP8A1*). ATP8A1 was shown to be elevated in post-mortem human juvenile hippocampal tissue from subjects with ASD, and the same study showed deficits in mouse sociability behavior due to induced *ATP8A1* overexpression [34]. However, *ATP8A1* knockout mice also showed differences in hippocampal function [34,35], suggesting that precise regulation of *ATP8A1* is necessary for normal hippocampal development and function. Three of twelve unaffected siblings also share this segment, so it predominantly is segregating to affected cases.

In family 10001, all eight affected family 10001 members share the segment on chromosome 3p21.31-p21.2 containing calcium voltage-gated channel auxiliary subunit alpha 2 delta 2 (*CACNA2D2*) and dedicator of cytokinesis 3 (*DOCK3*). Three of eight unaffected siblings also share this segment. Variants in *CACNA2D2* have been associated with childhood epilepsy [36,37] and opioid sensitivity [38]. Furthermore, ASD subjects have been identified with de novo functional mutations [39] or duplication events [40] in *CACNA2D2* (see supplemental material from both studies). Interestingly, *DOCK3* has also been associated with epilepsy [41] as well as attention deficit hyperactivity disorder [42]. According to ASD@Princeton, *DOCK3* is a known ASD gene, regulating post-synaptic density through stimulation of axonal outgrowth [43] and is a target of Fragile-X mental retardation protein [44]. An analysis of runs of homozygosity (ROH) in ASD cases identified several affected individuals with ROH in *DOCK3* [45]. Five affected members of family 8002 also share this segment.

Also in family 10001, seven affected, and one unaffected sibling, share the segment on chromosome 17q21.32 containing integrin subunit beta 3 (*ITGB3*), which has been previously associated with ASD [46–53], although not in a cohort of Irish ASD cases [54]. Considered by ASD@Princeton to be a known ASD gene, *ITGB3* plays a role in cell adhesion, and regulates serotonin levels through an interaction with the serotonin transporter [55]. Interestingly, the lone affected case that does not share this segment is the MZ twin. Complications arising during twin pregnancy and birth have been shown to increase the risk for ASD [56]; it is possible these non-genetic risks contributed significantly to this case.

In family 25002, all seven affected family 25002 members share the segment on chromosome 11p11.2 containing autophagy and beclin 1 regulator 1 (*AMBRA1*) and diacylglycerol kinase zeta (*DGKZ*), shared with six unaffected siblings, and the segment just downstream (11p11.2-p11.12) containing folate hydrolase 1 (*FOLH1*, discussed below), shared with seven unaffected siblings. *AMBRA1* functions in regulating neurogenesis, is prominently expressed in the striatum, hippocampus, and cortex [57,58], and AMBRA1 variations have been associated with schizophrenia [59,60]. Mouse models of *Ambra1* (+/-) heterozygous mutation show increased autism-like behavior exclusively among females coincided with lower protein expression among female vs. male heterozygous mice [58]. Likewise, variations in *DGKZ* were also linked to schizophrenia [59], and DGKZ is a target of FMRP [44].

Also in family 25002, the chromosome 15q25.1 segment, shared by six affected family 25002 members, and three of twelve unaffected siblings, contains 5,10-methenyltetrahydrofolate synthetase (*MTHFS*, discussed below) and aryl hydrocarbon receptor nuclear translocator 2 (*ARNT2*), which encodes a transcription factor involved in neurogenesis and plays a critical role in neurodevelopment of the hypothalamo-pituitary axis [61–63]. ARNT2 regulates transcription of oxygen-responsive genes by dimerizing with hypoxia-inducible factor 1 alpha and binding to promoter and enhancer segments [64,65]. Variation in *ARNT2* has been associated with ASD and schizophrenia [66].

Several other segments also showed evidence of overlap between all three families. There is a gene cluster of butyrophilin genes (including *BTN2A2* and *BTN3A3*) on chromosome 6p22 in another segment of interest. A component of human milk fat, butyrophilin glycoproteins are members of the immunoglobulin superfamily of transmembrane proteins [67] and are involved in lipid and sterol metabolism, and interact with HLA genes. They may be of interest given current hypotheses of a possible immune component to ASD susceptibility [68], particularly in regards to molecular mimicry as exposure to butyrophilin proteins can induce, as well as suppress, T-cell immune response to myelin oligodendrocyte glycoprotein (MOG) implicated in the pathogenesis of multiple sclerosis [69]. Of related interest, BTN2A2 has been associated with schizophrenia [70] and altered DNA methylation and expression of *BTN3A3* has been found in post-mortem brain tissue of individuals with bipolar disorder and schizophrenia [71]. Increased expression of *BTN2A2* was also found in postmortem brain tissue of individuals with schizophrenia [72].

ASD in these large, high-risk families is likely due to a combination of familial genetic variants, plus the interaction of each individual’s unique environment, that have a range of effects on phenotypic risk. For some cases it is possible that these risk factors also combine with unidentified functional *de novo* variation. This study highlighted segments of the genome that are shared between affected cases in three high-risk ASD families. It is promising that the most significant finding of the three individual families, the segment on 17q21.32 in family 10001, contained known ASD-risk gene *ITGB3*. Furthermore, both shared segments in family 25002 contained genes involved in folate metabolism, *FOLH1* and *MTHFS*. In the intestines, FOLH1 is required for folate absorption while in the brain, *FOLH1* is expressed by astrocytes, and facilitates the production of glutamate [73]. Variations in *FOLH1* have been associated with schizophrenia [74] and depression [75]. Activation of MTHFS intracellularly increases folate turnover and reduces folate concentrations [76]. A comparison of ASD cases and healthy controls found differences in the cerebrospinal fluid concentration of several folate metabolites [77], and folates have been explored or proposed as interventions in several neuropsychiatric disorders, including autism [77–79]. 5-methyltetrahydrofolate (5MTHF) crosses the blood-brain barrier and is the predominant form of folate in the central nervous system [80]. Although cerebrospinal 5MTHF concentrations have been shown to differ between ASD cases and healthy controls [77], another study did not find a correlation between 5MTHF and ASD symptoms in a cohort of young ASD cases sampled twice over a period of several years [81]. Central folate deficiency was identified among females with Rett Syndrome in a European study [82], though Neul et al (2005) failed to replicate this finding among a US sample of females with Rett Syndrome [83]. The invasive nature of cerebrospinal fluid collection and unpredictable nature of CSF 5MTHF level fluctuation, precludes routine CSF collection for 5MTHF levels during autism assessment [81].

Shared genomic segments analysis has the ability to narrow the search for meaningful risk variants that may be more common and have reduced penetrance, plus the capacity to tie variants to specific phenotypes in a cohort of affected cases. Furthermore, SGS is not limited to exome variation with a much smaller universe of potential variants. This study has demonstrated each of these points, and follow-up analyses will focus on identifying risk variants in these families using whole genome sequencing.

## Competing Interests Statement

All authors declare that there are no competing interests to disclose.

## Funding

This research was supported in part by R01 MH0994400, R01 CA163353, R01 CA134674, Simons Foundation Project 388196, the Margolis Foundation, and GCRC M01-RR025764 from the National Center for Research Resources. Partial support for all datasets within the Utah Population Database (UPDB) was provided by the University of Utah Huntsman Cancer Institute.

## Author Contributions

LJ maintained and prepared the biological samples. JM and DB provided expert clinical insight. NC, VR, RS, HC and TD developed, tested, and executed the SGS analysis. TD and HC wrote the manuscript, with input from all authors. All authors reviewed the manuscript.

## Acknowledgments

Additional computing resources were utilized through the Center for High Power Computing at the University of Utah (www.chpc.utah.edu). The analyses required extensive use of gnu parallel [84].

